# GPI-anchored Gas1 protein regulates cytosolic proteostasis in yeast

**DOI:** 10.1101/2023.05.26.542479

**Authors:** Yuhao Wang, Linhao Ruan, Rong Li

## Abstract

Decline in protein homeostasis (proteostasis) is a hallmark of cellular aging and aging-related diseases. Maintaining a balanced proteostasis requires a complex network of molecular machineries that govern protein synthesis, folding, localization, and degradation. Under proteotoxic stress, misfolded proteins that accumulate in cytosol can be imported into mitochondria for degradation via ‘mitochondrial as guardian in cytosol’ (MAGIC) pathway. Here we report an unexpected role of yeast Gas1, a cell wall-bound glycosylphosphatidylinositol (GPI)-anchored β-1,3-glucanosyltransferase, in differentially regulating MAGIC and ubiquitin-proteasome system (UPS). Deletion of Gas1 inhibits MAGIC but elevates polyubiquitination and UPS-mediated protein degradation. Interestingly, we found that Gas1 exhibits mitochondrial localization attributed to its C-terminal GPI anchor signal. But this mitochondria-associated GPI anchor signal is not required for mitochondrial import and degradation of misfolded proteins via MAGIC. By contrast, catalytic inactivation of Gas1 via the *gas1^E161Q^*mutation inhibits MAGIC but not its mitochondrial localization. These data suggest that the glucanosyltransferase activity of Gas1 is important for regulating cytosolic proteostasis.

## Introduction

Many aging-related degenerative diseases are associated with loss of proteostasis characterized by protein misfolding and formation of protein aggregates (1–3). Major protein quality control pathways such as UPS and autophagy play essential roles in clearance and turnover of misfolded proteins (4). Mitochondria are key organelles not only in cellular metabolism (5) but also in clearing misfolded proteins via a process termed as ‘mitochondrial as guardian in cytosol’ (MAGIC) (6–8). In budding yeast, misfolded cytosolic proteins can translocate into mitochondria through mitochondrial import channels (6). The yeast Lon protease, Pim1, is involved in the degradation of imported misfolded proteins in the mitochondrial matrix, which in turn facilitates the dissolution of cytosolic protein aggregates associated with mitochondria (6,7). Likewise, a model unstable cytosolic protein (FlucDM) can be imported into the mitochondrial matrix of human RPE-1 cells (6). Proteasomal inhibition by MG132 in HeLa cells can induce the import of unfolded cytosolic GFP into mitochondria through the joint action of mitochondrial outer membrane protein FUNDC1 and cytosolic chaperone HSC70 (9). Many disease-related proteins such as α-synuclein, FUS, and TDP-43 have been shown to accumulate in mitochondria of various human cell models and influence mitochondrial fitness (10–12). These findings suggest that MAGIC may be a conserved process that connects cytosolic proteostasis with mitochondrial functions.

To elucidate the molecular mechanism and regulation of MAGIC in yeast, we conducted an unbiased genetic screen using the non-essential yeast knockout library that unveiled potential regulators of MAGIC including Gas1 (7). Gas1 (glycophospholipid-anchored surface protein 1) is a β-1,3-glucanosyltransferase anchored to the outer leaflet of the plasma membrane via a GPI moiety and crosslinked with the cell wall (13–16). Gas1 elongates the β-1,3-glucan chains and plays an important role in expansion and remodeling of the cell wall (17). Here, we show that Gas1 is a positive regulator of MAGIC, and loss of Gas1 prevents the accumulation and degradation of misfolded proteins in mitochondria. We also provide insights into the subcellular localization of Gas1-GFP by showing that its C-terminal GPI anchor signal, but not the main part of the protein, localizes to mitochondria. Despite this atypical localization, the catalytic activity of Gas1, but not the mitochondria-associated GPI anchor signal, is required for maintaining MAGIC. Furthermore, two other cell wall mutants show moderate inhibition of MAGIC, suggesting that maintenance of cell wall integrity in general may be important for MAGIC. In addition to altered MAGIC phenotypes, Gas1-deficient cells exhibit elevated protein polyubiquitination and UPS-mediated degradation of an N-end rule substrate (Ub-R-EGFP). Taken together, our work identifies an unexpected role of Gas1 in regulating proteostasis pathways and suggests an underappreciated connection between cytosolic proteostasis and cell wall integrity.

## Results

### Gas1 is required for mitochondrial accumulation and degradation of misfolded proteins

We conducted an unbiased genetic screen and subsequent validations that uncovered potential MAGIC regulators in yeast by performing the split-GFP assay in the yeast non-essential knockout library (6,7). In brief, cytosolic proteins such as Lsg1, an endogenous aggregation-prone protein (6), or FlucSM, a destabilized firefly luciferase mutant and a sensor of the proteostasis stress (6,7,18), were tagged with the eleventh β-strand of GFP (GFP_11_), and the first ten β-strands of GFP (GFP_1-10_) were targeted to mitochondrial matrix by fusing with the mitochondrial matrix protein Grx5 (Grx5-GFP_1-10_) or the mitochondrial targeting sequence of Subunit 9 of mitochondrial ATPase (Su9 MTS) from *Neurospora crassa* (MTS-mCherry-GFP_1-10_). In wild-type cells, proteotoxic stresses such as heat shock (6) or acute overexpression of misfolded proteins induced by estradiol (7) resulted in an increased GFP fluorescence (spGFP signal) within mitochondria (Figure 1A-C). We analyzed each mutant before and after stress and identified gene deletions that gave rise to differential spGFP patterns compared to the wild-type cell (7).

**Figure 1.**
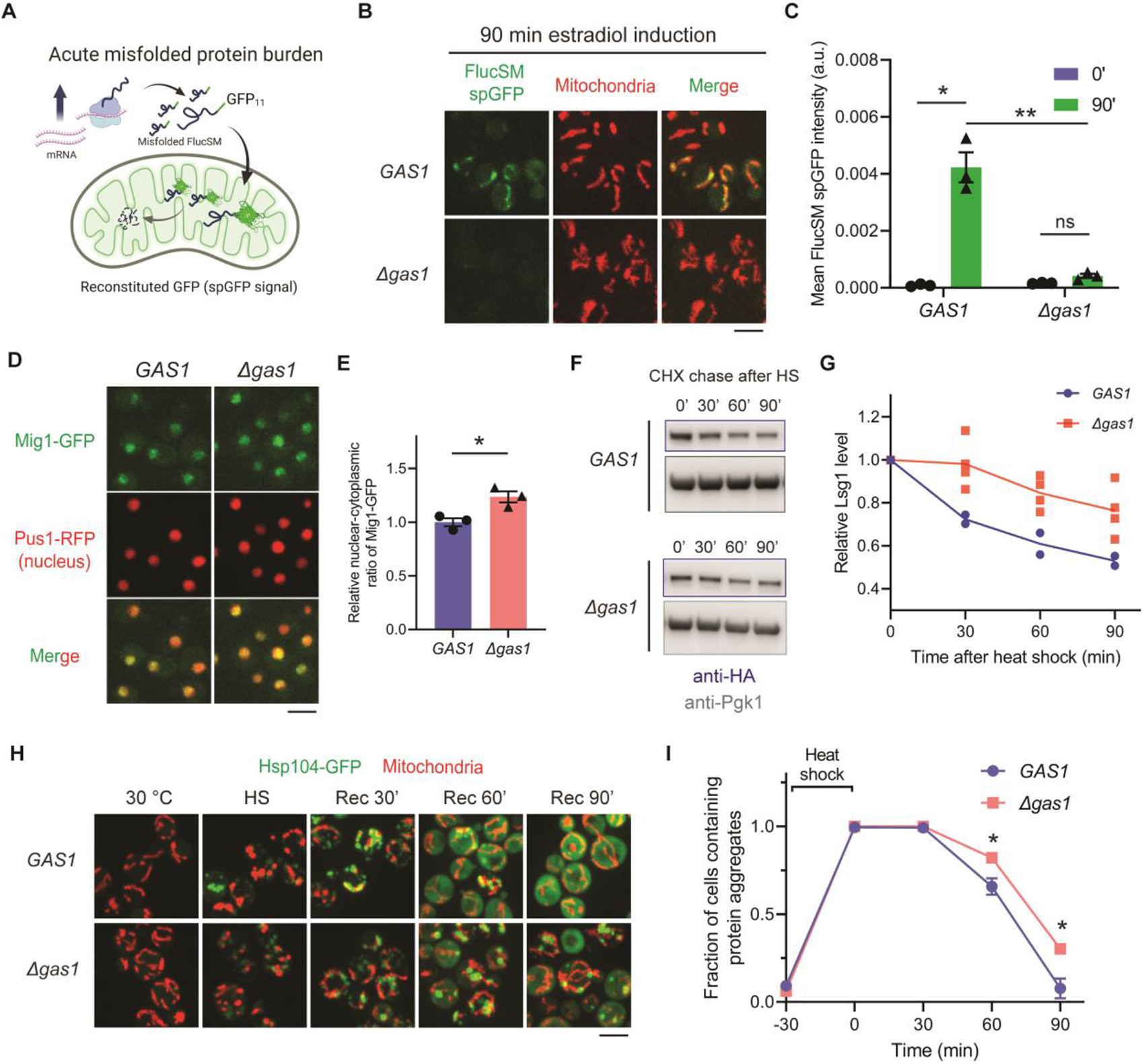
Loss of Gas1 inhibits mitochondrial accumulation and degradation of misfolded proteins. (**A**) Schematic diagram of FlucSM split-GFP assay upon proteotoxic stress associated with elevated misfolded protein burden. (**B**, **C**) Images (B) and quantification (C) of FlucSM spGFP signal in mitochondria of wild-type *GAS1* and *Δgas1* cells after estradiol induction. Shown in (C): means ± SEM of FlucSM spGFP intensities with arbitrary unit (a.u.). Paired two-tailed *t*-test between no induction (0’) and 90 min induction (90’). Unpaired two-tailed *t*-test between *GAS1* and *Δgas1* cells after 90 min induction. Three biological repeats. (**D**, **E**) Images (D) and quantification (E) of the nuclear-cytoplasmic translocation of Mig1-GFP signal. Shown in (E): means ± SEM of normalized Mig-GFP nuclear-cytoplasmic ratios. Unpaired two-tailed *t*-test. (**F**, **G**) Immunoblots (F) and quantifications (G) showing data points and means of Lsg1-HA degradation *in vivo* after heat shock (HS) in the presence of cycloheximide (CHX). (**H, I**) Dissolution of protein aggregates labeled with Hsp104-GFP in *GAS1* and *Δgas1* cells during recovery (Rec) after heat shock. Shown in (I): means ± SEM of fractions of cells containing protein aggregates. Unpaired two-tailed *t*-test. \**P* < 0.05; \*\**P* < 0.01; \*\*\**P* < 0.001; ns, not significant, *P* > 0.05. Scale bars, 5 μm.

*Δgas1* was one of mutants that failed to show an increased FlucSM spGFP signal after stress compared to the wild-type control (Figure 1B and C) (7). Because activation of Snf1 kinase, the yeast homolog of human AMP-activated protein kinase, inhibits the import of misfolded proteins into mitochondria (7), we first asked if Snf1 activity is elevated in *Δgas1* by using the nuclear-cytoplasmic transport of Mig1-GFP as a reporter. Under glucose restriction, Mig1 is phosphorylated by active Snf1 and exported from the nucleus (7,19). However, Mig1-GFP signal remained in the nucleus in *Δgas1* mutant cells and even exhibited slightly more enrichment compared to the control (Figure 1D and E). This data argues against the possibility that the inhibition of MAGIC in *Δgas1* mutant was primarily caused by Snf1 activation. In addition to the reduced FlucSM spGFP signal, we used immunoblots to assess the degradation of misfolded proteins, and performed time-lapse imaging to track the disaggregation of cytosolic protein aggregates, both of which have been shown to depend on MAGIC (6). We found that the post-heat shock degradation of Lsg1 was significantly delayed in *Δgas1* (Figure 1F and G). The dissolution of protein aggregates labeled with protein disaggregase Hsp104-GFP after heat shock was also impaired, even though the association of aggregates with mitochondria appeared unaffected (Figure 1H and I). In sum, these data suggest that Gas1 is a regulator of the mitochondria-mediated pathway for degrading certain cytosolic aggregation-prone proteins.

### Re-analysis of the intracellular localization of Gas1-GFP reveals an unexpected mitochondrial targeting domain

Gas1 was previously shown to localize to mitochondria in addition to cell periphery and ER (20,21). It was therefore reasonable for us to hypothesize that the mitochondrial localization of Gas1 is related to its role in MAGIC. To test this, it was necessary to gain an understanding of how this purported cell wall protein is targeted to mitochondria. By using confocal microscopy to observe the subcellular localization of the C-terminally tagged Gas1-GFP, we confirmed that a portion of GFP fluorescence colocalized with mitochondria labeled with MTS-mCherry, while the other fraction of GFP fluorescence appeared to be in the ER and nuclear periphery (Figure 2A), which was reported previously (21). By using super-resolution microscopy, we found that mitochondrial Gas1-GFP signal delineates the contour of the mitochondrial matrix, suggesting its localization in the mitochondrial membranes or intermembrane space (Figure 2B). Importantly, our Gas1-GFP strain was resistant to treatment with calcofluor white (CFW), a cell wall destabilizing reagent (22), whereas *Δgas1* mutant with cell wall defects failed to grow on CFW-containing plates (Supplementary Figure 1A). This data suggests that C-terminal GFP tagging of Gas1 preserves its function in maintaining cell wall integrity.

**Figure 2.**
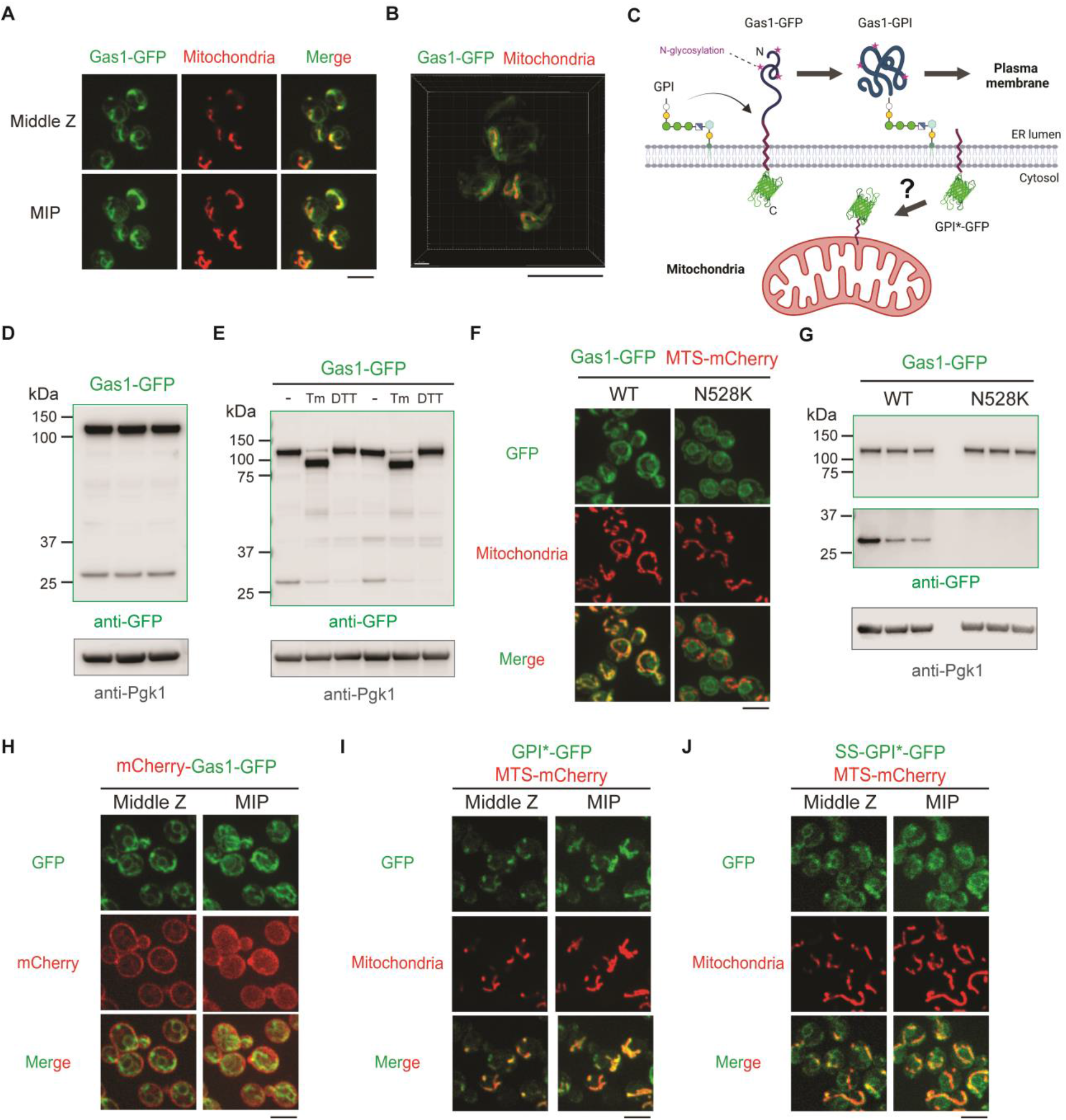
C-terminal GPI anchor signal of Gas1 localizes to mitochondria. (**A**) Images of Gas1-GFP, showing partial colocalization with mitochondrial matrix labeled by MTS-mCherry. The ring-like structure may represent ER/nuclear periphery as reported previously (21). MIP, maximal intensity projection. (**B**) 3D rendering of super-resolution images of Gas1-GFP, showing GFP signals outside the mitochondrial matrix labeled with MTS-mCherry. (**C**) Working model of GPI addition and peptide cleavage during Gas1 maturation in the ER. (**D**) Immunoblots of whole cell lysates showing two major species with GFP. Three biological repeats. (**E**) Immunoblots of whole cell lysates after tunicamycin (Tm) or DTT treatment. (**F, G**) Images (F) and immunoblots (G) of whole cell lysates expressing Gas1-GFP or Gas1^N528K^-GFP. (**H**-**J**) Images of cells expressing an ectopic copy of mCherry-Gas1-GFP (H), GPI*-GFP (I), or SS-GPI*-GFP (J). MIP, maximal intensity projection. Scale bars, 5 μm.

The trafficking and post-translational modification of Gas1 are intricately coupled processes (13-15,23,24). To reach the cell surface, newly synthesized Gas1 protein is targeted to ER by a canonical N-terminal signal sequence (SS) (13–15). A cleavable hydrophobic sequence at the C terminus temporarily anchors Gas1 on the ER membrane and serves as a GPI anchor signal (GPI*) that is necessary for the addition of a GPI moiety via the GPI-anchor transamidase complex (13,15,23). As a result, we conceived that the cleaved GPI* should bear the GFP tag (GPI*-GFP), and the GPI-anchored Gas1 (Gas1-GPI) is transported to plasma membrane through the secretory pathway where it is progressively modified by N-glycosylation and O-glycosylation in the ER and Golgi (13,23) (Figure 2C). In line with this model, anti-GFP immunoblot analysis on the yeast whole cell lysate of Gas1-GFP strain showed two major species (Figure 2D). A higher molecular weight (MW) species exhibited the expected size of N-glycosylated uncleaved Gas1-GFP (∼120 kDa), as confirmed by treatment with tunicamycin, an inhibitor of N-acetylglucosamine transferases (25), which decreased its MW to the unmodified Gas1-GFP (∼90 kDa) (Figure 2D and E). By contrast, dithiothreitol (DTT) treatment, which reduces disulfide bonds of ER proteins and causes the unfolded protein stress (26), had no effect on N-glycosylation (Figure 2E). The low MW species appeared to be comparable to GPI*-GFP (∼28 kDa) (Figure 2D). Interestingly, either treatment with tunicamycin or DTT reduced the amount of low MW species, suggesting that ER stress may compromise the cleavage of GPI* regardless of the presence of N-glycosylation (Figure 2E).

We next tested if the cleavage of GPI* is responsible for the mitochondrial localization of Gas1. The amino acid sequence at and adjacent to the GPI anchor attachment site of Gas1 (N528) is important for efficient peptide cleavage (24) (Supplementary Figure 1B). Mutation of N528 to a lysine residue (N528K) abolishes the glycolipid transfer and peptide cleavage, and as a result, Gas1 remains unprocessed and only N-glycosylated in the ER (24). We found that Gas1^N528K^-GFP failed to exhibit any mitochondrial localization (Figure 2F), and as expected the low MW species also disappeared in the anti-GFP immunoblot (Figure 2G). This result indicates that mitochondrial GFP fluorescence requires the cleavage of GPI*.

To differentially observe the N and C terminal portion of Gas1 after GPI* cleavage, we fused the Gas1 protein with a mCherry at its N terminus after the ER-targeting signal peptide, and simultaneously with a GFP at its C terminus following the GPI anchor signal (mCherry-Gas1-GFP). The mCherry signal was mostly detected on the cell periphery where the GFP fluorescence was largely absent, whereas only GFP but not mCherry signal was detected on mitochondria (Figure 2H). Only at the ER/nuclear periphery were both mCherry and GFP colocalized, which likely represented the unprocessed Gas1 (Figure 2H). This dual-color pattern further confirms that after GPI* cleavage, the N-terminal main portion of the protein is trafficked to the cell periphery whereas GPI* is targeted to mitochondria.

To determine if GPI* is sufficient for mitochondrial localization, we ectopically expressed the C-terminally tagged GPI*-GFP with or without an additional N-terminal ER-targeting signal sequence (SS-GPI*-GFP) under a native *GAS1* promoter. GPI*-GFP signal was predominantly localized to mitochondria, whereas SS-GPI*-GFP showed both ER and mitochondrial localization (Figure 2I and J). This data suggests that the GPI anchor signal of Gas1 could be a mitochondrial targeting sequence even in the presence of an ER-targeting sequence.

### Enzymatic activity of Gas1 rather than its GPI anchor signal is required for MAGIC

We next asked if the mitochondrial localized GPI anchor signal of Gas1 is required for maintaining MAGIC, a process dependent on functional mitochondrial import (6). To avoid affecting the targeting to plasma membrane and function of Gas1 in the cell wall, we replaced the endogenous GPI anchor signal of Gas1 by that of Gas3 and Gas5, two closely related Gas family glucanosyltransferases with C-terminal GPI anchor signals (Supplementary Figure 1B) (15). Yeast strains expressing recombinant proteins (Gas1-GPI*_Gas3_ and Gas1-GPI*_Gas5_) were resistant to treatment with CFW, and these proteins were fully glycosylated at a level comparable to the wild-type Gas1 (Supplementary Figure 1C and D), indicating normal processing and cell wall function of the recombinant Gas1. Importantly, by C-terminal tagging with GFP, neither recombinant protein exhibited mitochondrial localization: Gas1-GPI*_Gas3_-GFP was largely confined to the ER, and Gas1-GPI*_Gas5_-GFP signal appeared to be modestly present in the vacuole (Figure 3A). Mitochondrial accumulation of misfolded FlucSM after acute overexpression was not affected by GPI* replacement (Figure 3B and C). Misfolded endogenous Lsg1 was also degraded normally after heat shock (Figure 3D and E). These results suggest that the GPI anchor signal of Gas1 and its mitochondrial localization are not involved in the regulation of MAGIC.

**Figure 3.**
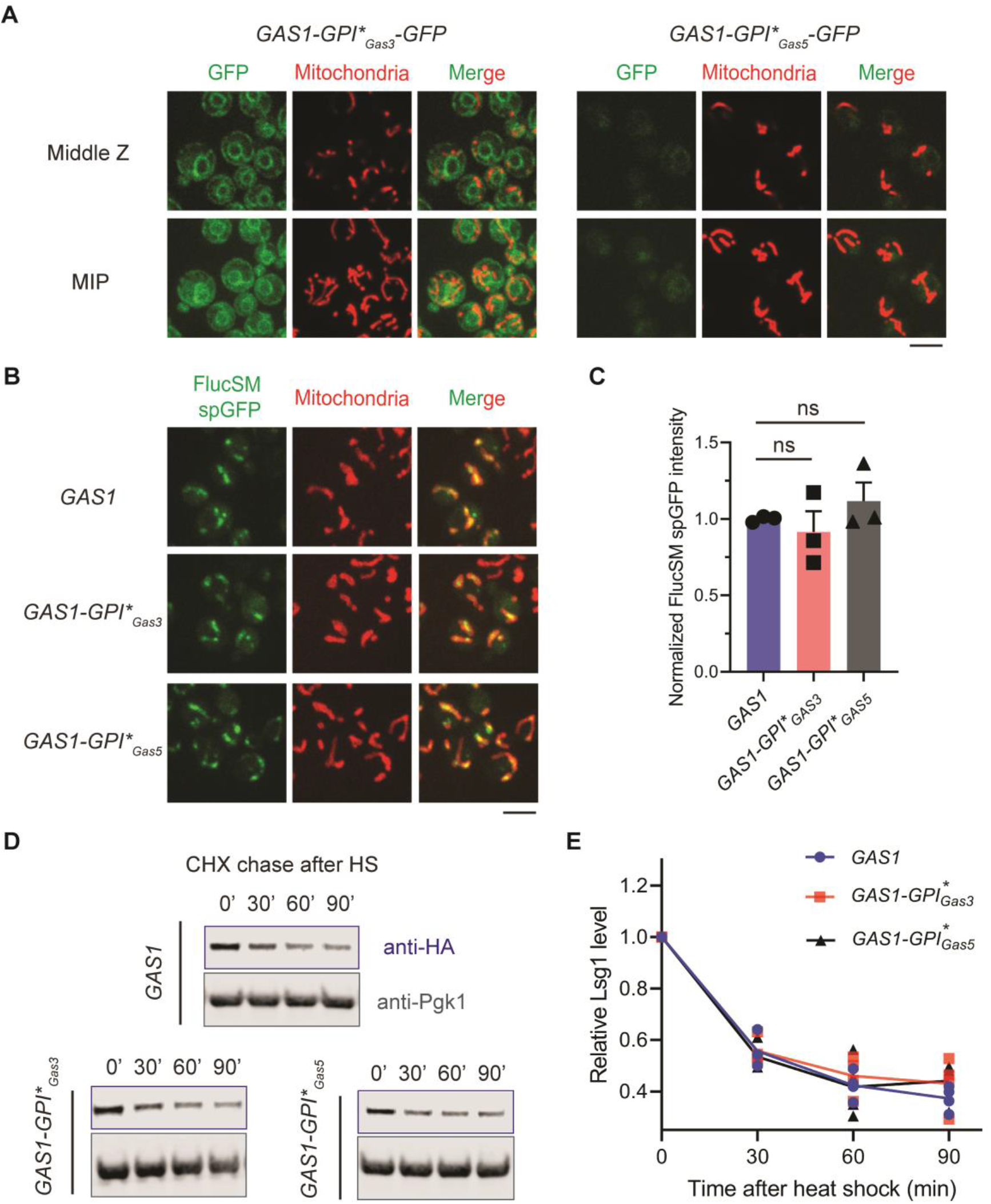
Mitochondria-associated GPI anchor signal of Gas1 is not required for MAGIC. (**A**) Images of cells expressing an ectopic copy of Gas1-GPI*_Gas3_-GFP or Gas1-GPI*_Gas5_-GFP and MTS-mCherry labelling mitochondria. MIP, maximal intensity projection. (**B, C**) Images (B) and quantification (C) of FlucSM spGFP signal in mitochondria of cells expressing wild-type or recombinant Gas1 with GPI* replaced. Shown in (C): means ± SEM of normalized FlucSM spGFP intensities. ns, not significant, *P* > 0.05. Unpaired two-tailed *t*-test. (**D**, **E**) Immunoblots (D) and quantifications (E) of Lsg1-HA degradation *in vivo* after heat shock (HS) in the presence of cycloheximide (CHX). Scale bars, 5 μm.

We therefore tested if the glucanosyltransferase activity of Gas1 is required for MAGIC by studying the effect of the catalytically inactive *gas1^E161Q^*mutation (21,27), which also caused CFW sensitivity (Supplementary Figure 1E). Like the wild-type protein, Gas1^E161Q^ was fully glycosylated and likely trafficked to the cell surface (Supplementary Figure 1F). Gas1^E161Q^-GFP showed both ER and mitochondrial GFP localization, indicative of normal cleavage of GPI* (Figure 4A). But like *Δgas1*, this mutant exhibited defects in proteostasis and MAGIC, including delayed dissolution of protein aggregates after heat shock and reduced accumulation of misfolded FlucSM in mitochondria (Figure 4B-E). These results suggest that the enzymatic activity of Gas1 is essential for MAGIC.

**Figure 4.**
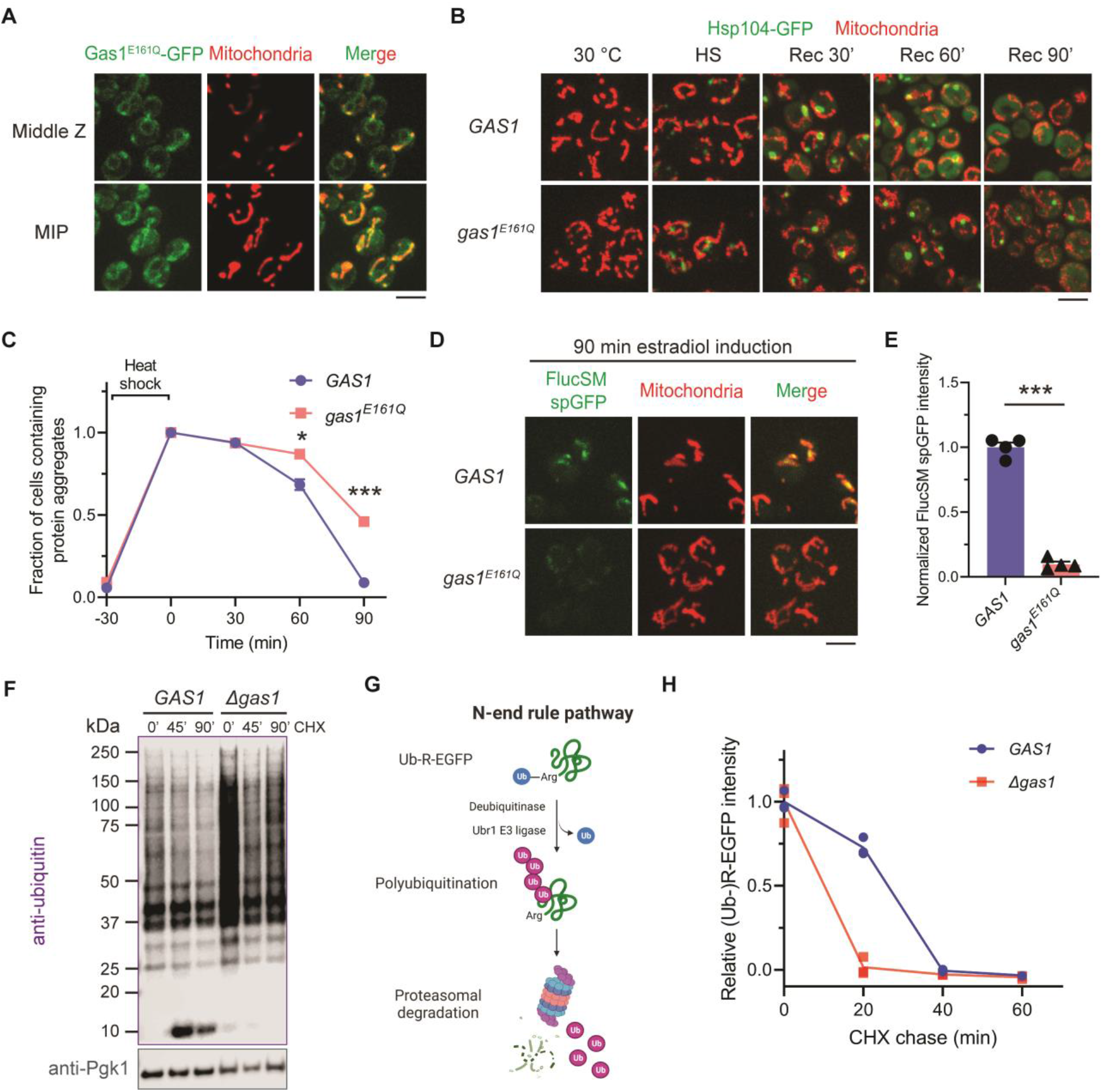
Loss of Gas1 activity inhibits MAGIC but elevates UPS-mediated degradation. (**A**) Images of cells expressing an ectopic copy of Gas1^E161Q^-GFP and MTS-mCherry. MIP, maximal intensity projection. (**B**, **C**) Dissolution of protein aggregates labeled with Hsp104-GFP in *GAS1* and *gas1^E161Q^* cells after heat shock. Unpaired two-tailed *t*-test. (**D**, **E**) Images (D) and quantification (E) of FlucSM spGFP in mitochondria of *GAS1* and *gas1^E161Q^* cells after estradiol treatment for 90 min. Shown in (E): means ± SEM of normalized FlucSM spGFP intensities. Unpaired two-tailed *t*-test. (**F**) Anti-ubiquitin immunoblots of cell lysates before and after cycloheximide (CHX) treatment, showing an accumulation of polyubiquitinated species with high molecular weight in *Δgas1* cells. (**G**) Diagram of N-end rule pathway using Ub-R-EGFP as a proteasomal substrate. (**H**) Degradation of copper-induced Ub-R-EGFP signal *in vivo* in wild-type and *Δgas1* cells. Three biological repeats. \**P* < 0.05; \*\*\**P* < 0.001. Scale bars, 5 μm.

To determine whether loss of cell wall integrity could inhibit MAGIC in general, we examined two other cell wall mutants with the deletion of *BGL2* encoding endo-β-1,3-glucanase Bgl2 (28), or *GAS5* encoding another β-1,3-glucanosyltransferase (14,15). Both mutants showed sensitivity to CFW, albeit much milder than *Δgas1* (Supplementary Figure 1G). They also caused a moderate reduction in mitochondrial accumulation of FlucSM in spGFP assay (Supplementary Figure 1H), which was less severe than Gas1-deficient cells (*Δgas1* in Figure 1C; *gas1^E161Q^* in Figure 4E). This analysis indicates that cell wall maintenance is indeed important for MAGIC.

### Effect of Gas1 deficiency on UPS

To gain insights into whether Gas1 plays a broader role in other proteostasis pathways, we examined the effect of *Δgas1* on UPS. Surprisingly, anti-ubiquitin immunoblot analysis on the whole cell lysate of the mutant revealed a drastic increase in overall protein polyubiquitination compared to the wild-type cell (Figure 4F). Over-accumulated polyubiquitinated proteins gradually disappeared after inhibiting protein synthesis by cycloheximide (Figure 4F). In contrast to the delayed degradation of MAGIC substrate, loss of Gas1 resulted in a faster degradation of non-misfolded Ub-R-EGFP, an N-end rule substrate of the ubiquitin-proteosome system (Figure 4G and H) (29–31). In summary, our data suggests that loss of Gas1 activity may enhance UPS while inhibiting MAGIC.

## Discussion

Yeast Gas1 is a GPI-anchored β-1,3-glucanosyltransferase that elongates the β-1,3-glucan chains and plays a critical role in the dynamic remodeling of the cell wall (14,15). The yeast null mutant (*Δgas1*) shows cell wall defects characterized by round cells, CFW sensitivity, reduced growth rate, and decreased cell viability during the stationary phase (22,32,33). Gas1 deficiency also triggers the cell wall integrity signaling pathway and compensatory responses such as an increase in chitin synthesis and deposition to rescue cell wall integrity (32,34). In addition to the established function at the cell wall, Gas1 was shown to play roles in intracellular processes such as DNA damage response (35), locus-specific transcriptional silencing (21), and ER stress response (36). Deletion of Gas1 increases cellular sensitivity to DNA damages caused by genotoxins (35), and loss of its enzymatic activity leads to defective telomeric silencing and elevated rDNA silencing (21). Because C-terminally tagged Gas1-GFP shows intracellular localizations in ER/nuclear periphery (as well as mitochondria) (20,21), it has been proposed that Gas1 is involved in post-translational glycosylation of putative chromatin components (21,35). Loss of Gas1 also elevates the unfolded protein response (UPR) pathway and renders the mutant cells resistant to tunicamycin-induced ER proteotoxic stress (36). Both UPR and ER-associated degradation (ERAD) maintain proteostasis in the ER and may in turn promote the secretion and glycosylation of essential plasma membrane and cell wall proteins (37). However, detailed mechanisms underlying the intracellular functions of Gas1 remain unclear.

Mitochondria are important organelles in cellular metabolism and cytosolic proteostasis (5–9,38). Loss of cytosolic proteostasis is manifested by protein misfolding and formation of protein aggregates, which are often tethered to intracellular organelles such as ER and mitochondria (39,40). Under proteotoxic stress, certain misfolded cytosolic proteins can be translocated into and degraded inside mitochondria through MAGIC pathway (6). Our data suggests that Gas1 function is important for cytosolic proteostasis mechanisms including MAGIC and UPS. Specifically, Gas1 deficiency inhibits the mitochondrial accumulation and degradation of aggregation-prone proteins but promotes UPS-mediated degradation of a non-misfolded N-end rule substrate. One caveat of this study is that most of our observations were based on a few model substrates such as FlucSM, Lsg1, and Ub-R-EGFP. Thus the generality and the underlying mechanisms of our observations are presently unknown.

While investigating if mitochondrial targeting is important for the role of Gas1 in MAGIC, we unexpectedly identified the C-terminal GPI anchor signal of Gas1 (GPI*_Gas1_) as an unconventional mitochondrial targeting sequence. GPI*_Gas1_ does not bear typical amphiphilic helices like Su9 MTS from *Neurospora crassa* but likely contains a transmembrane domain, which was not present in GPI*_Gas3_ or GPI*_Gas5_ (Supplementary Figure 2A). It is worth noting that a previous study showed that GPI*_Gas1_ with an N-terminal GFP tag (also denoted as GFP-Gas1GPI) only exhibits ER localization (41). We speculate that a well-folded GFP at the N terminus may hinder the import or association of GPI* with mitochondria. GPI* may have a dual targeting specificity that allows its route to ER when mitochondrial targeting is impaired. Anti-GPI*_Gas1_ immunoblots on mitochondrial fraction or immunofluorescence staining may be necessary to further validate its subcellular localization. Also, it will be interesting to test if some other GPI-anchored proteins in yeast or higher organisms contain a GPI anchor signal with similar hydrophobic properties and mitochondrial targeting capacity.

By replacing the GPI* of Gas1 by that of Gas3 or Gas5, we showed that this mitochondria-associated peptide is dispensable for mitochondrial accumulation of misfolded FlucSM and post-heat shock degradation of Lsg1 by MAGIC. By contrast, the catalytically inactive Gas1^E161Q^ protein exhibited normal maturation and mitochondrial localization, but still failed to maintain MAGIC. Importantly, two other cell wall mutants with moderate CFW sensitivity also displayed mild inhibition of MAGIC, suggesting that cell wall integrity may be important for MAGIC. The stronger phenotypes of *Δgas1* or *gas1^E161Q^* mutant than others in terms of CFW sensitivity and MAGIC inhibition may be attributed to either more severe cell wall defects or Gas1-specific effects that remain to be determined. One possibility is that loss of cell wall associated Gas1 activity may trigger the downstream signaling pathways that separately or jointly regulate MAGIC and UPS. Our results, however, do not rule out the possibility that the intracellular fraction of Gas1 in the ER/nuclear periphery plays a more direct role in proteostasis.

## Materials and Methods

### Yeast strains and growth conditions

Yeast strains used in this study are based on the BY4741 strain background and listed in Table 1. Gene deletion and protein tagging were performed by using PCR-mediated homologous recombination and verified by PCR genotyping. *GAS1-GFP* and *MIG1-GFP* strain were obtained from the yeast GFP clone collection (20). *PUS1-RFP* strain was retrieved from yeast RFP library (42). *Δbgl2* and *Δgas5* were collected from the non-essential yeast knockout library (43). For the FlucSM split-GFP assay, GFP_1-10_ was fused with the mitochondrial matrix protein Grx5 under *GPD* promoter and stably integrated into yeast genome at the *TRP1* locus. FlucSM-HA-GFP_11_ (6) under *GAL1* promoter and GEM transcriptional factor (44) were cloned and stably integrated into yeast genome. Mitochondria were labeled with Tom70-mCherry in the split-GFP assay or with MTS-mCherry in other imaging experiments.

**Table 1.**
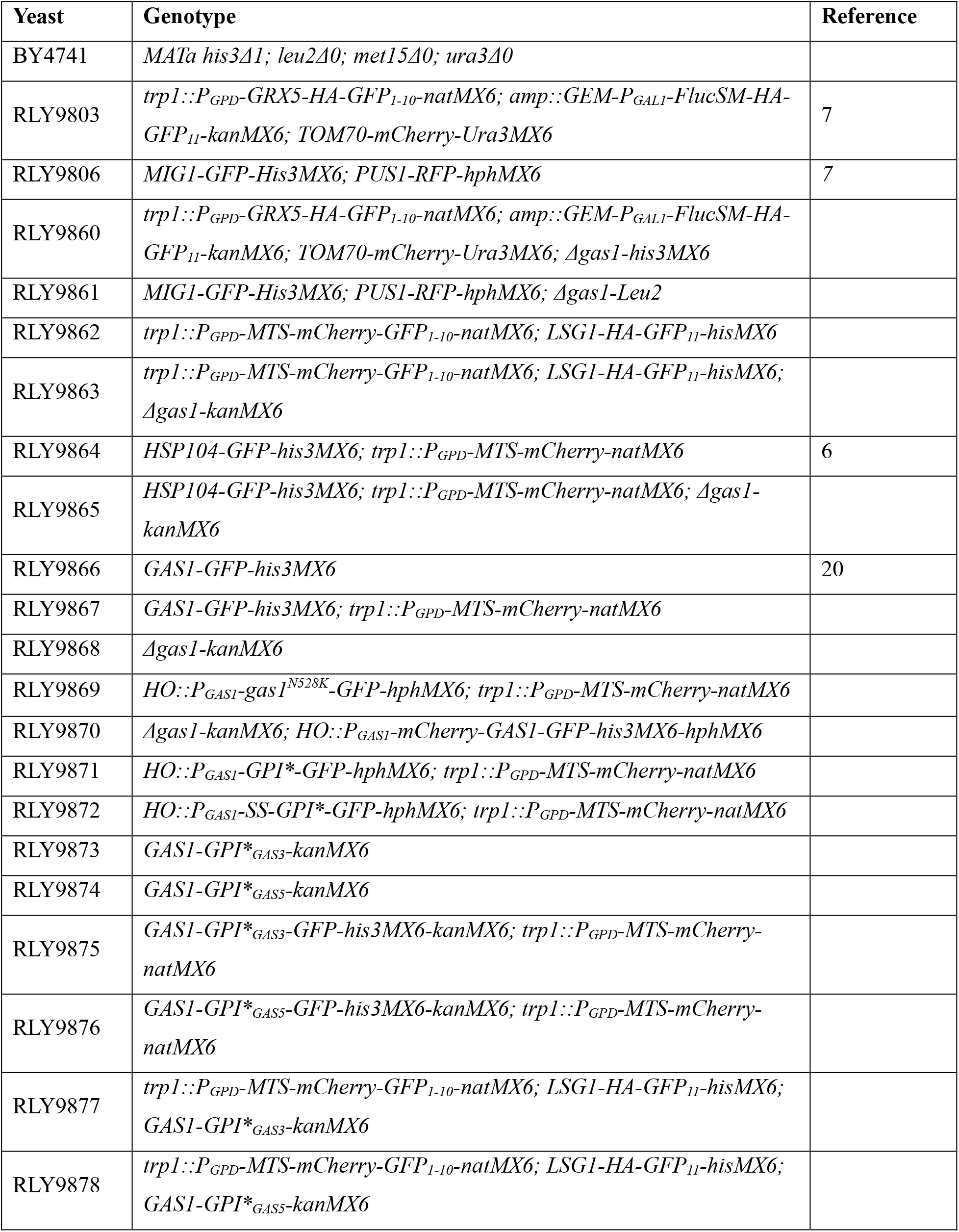
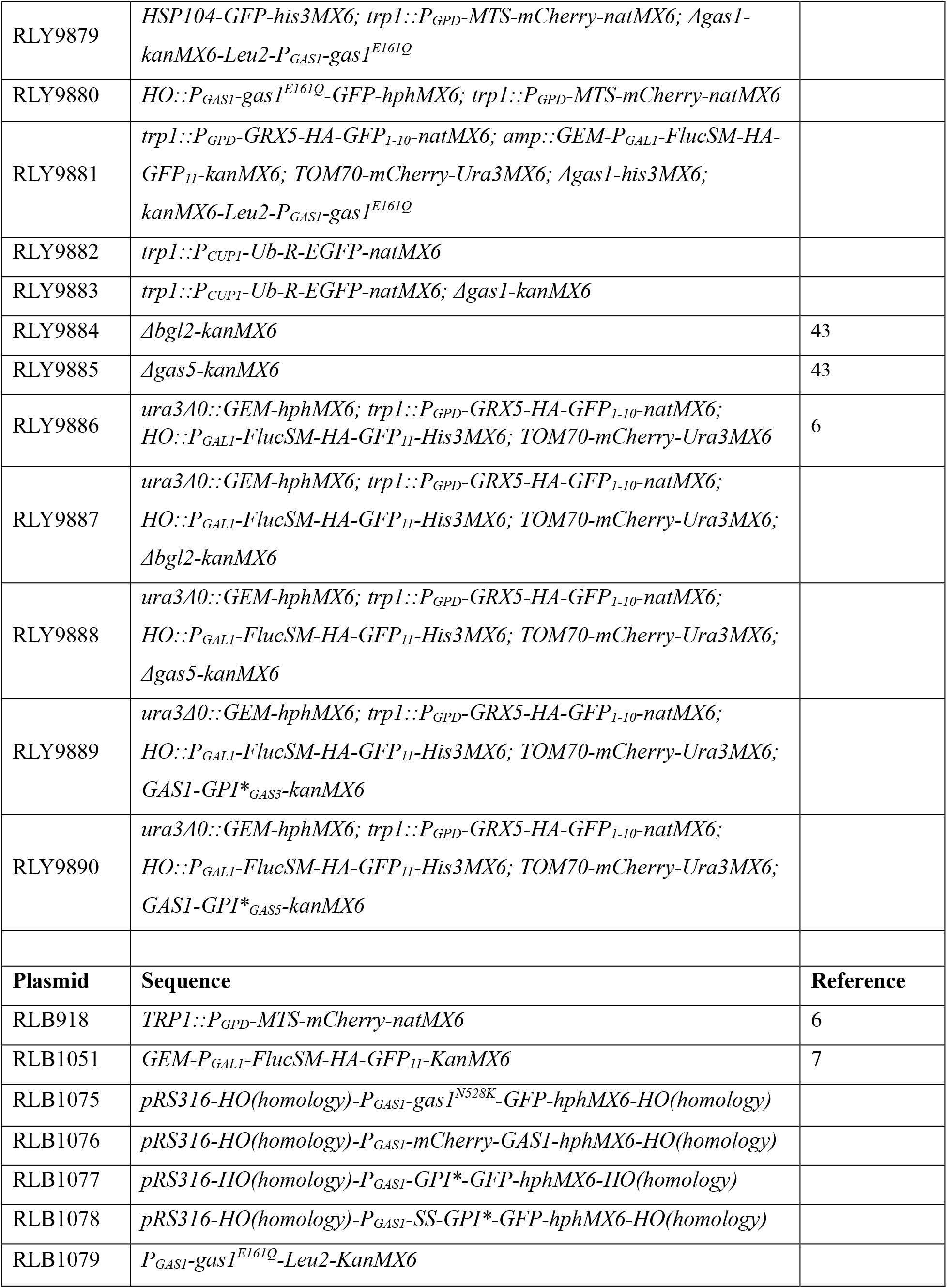
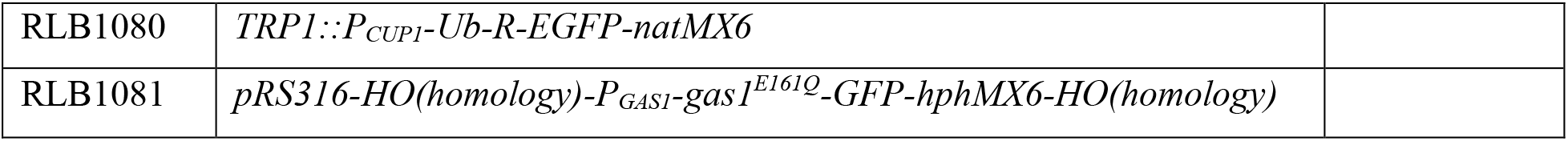
List of yeast strains and plasmids.

| Yeast | Genotype | Reference |
| --- | --- | --- |
| BY4741 | *MATa his3Δ1; leu2Δ0; met15Δ0; ura3Δ0* |  |
| RLY9803 | *trp1::P<sub>GPD</sub>-GRX5-HA-GFP<sub>1-10</sub>-natMX6; amp::GEM-P<sub>GAL1</sub>-FlucSM-HA-GFP<sub>11</sub>-kanMX6; TOM70-mCherry-Ura3MX6* | 7 |
| RLY9806 | *MIG1-GFP-His3MX6; PUS1-RFP-hphMX6* | 7 |
| RLY9860 | *trp1::P<sub>GPD</sub>-GRX5-HA-GFP<sub>1-10</sub>-natMX6; amp::GEM-P<sub>GAL1</sub>-FlucSM-HA-GFP<sub>11</sub>-kanMX6; TOM70-mCherry-Ura3MX6; Δgas1-his3MX6* |  |
| RLY9861 | *MIG1-GFP-His3MX6; PUS1-RFP-hphMX6; Δgas1-Leu2* |  |
| RLY9862 | *trp1::P<sub>GPD</sub>-MTS-mCherry-GFP<sub>1-10</sub>-natMX6; LSG1-HA-GFP<sub>11</sub>-hisMX6* |  |
| RLY9863 | *trp1::P<sub>GPD</sub>-MTS-mCherry-GFP<sub>1-10</sub>-natMX6; LSG1-HA-GFP<sub>11</sub>-hisMX6; Δgas1-kanMX6* |  |
| RLY9864 | *HSP104-GFP-his3MX6; trp1::P<sub>GPD</sub>-MTS-mCherry-natMX6* | 6 |
| RLY9865 | *HSP104-GFP-his3MX6; trp1::P<sub>GPD</sub>-MTS-mCherry-natMX6; Δgas1-kanMX6* |  |
| RLY9866 | *GAS1-GFP-his3MX6* | 20 |
| RLY9867 | *GAS1-GFP-his3MX6; trp1::P<sub>GPD</sub>-MTS-mCherry-natMX6* |  |
| RLY9868 | *Δgas1-kanMX6* |  |
| RLY9869 | *HO::P<sub>GAS1</sub>-gas1<sup>N528K</sup>-GFP-hphMX6; trp1::P<sub>GPD</sub>-MTS-mCherry-natMX6* |  |
| RLY9870 | *Δgas1-kanMX6; HO::P<sub>GAS1</sub>-mCherry-GAS1-GFP-his3MX6-hphMX6* |  |
| RLY9871 | *HO::P<sub>GAS1</sub>-GPI\*-GFP-hphMX6; trp1::P<sub>GPD</sub>-MTS-mCherry-natMX6* |  |
| RLY9872 | *HO::P<sub>GAS1</sub>-SS-GPI\*-GFP-hphMX6; trp1::P<sub>GPD</sub>-MTS-mCherry-natMX6* |  |
| RLY9873 | *GAS1-GPI\*<sub>GAS3</sub>-kanMX6* |  |
| RLY9874 | *GAS1-GPI\*<sub>GAS5</sub>-kanMX6* |  |
| RLY9875 | *GAS1-GPI\*<sub>GAS3</sub>-GFP-his3MX6-kanMX6; trp1::P<sub>GPD</sub>-MTS-mCherry-natMX6* |  |
| RLY9876 | *GAS1-GPI\*<sub>GAS5</sub>-GFP-his3MX6-kanMX6; trp1::P<sub>GPD</sub>-MTS-mCherry-natMX6* |  |
| RLY9877 | *trp1::P<sub>GPD</sub>-MTS-mCherry-GFP<sub>1-10</sub>-natMX6; LSG1-HA-GFP<sub>11</sub>-hisMX6; GAS1-GPI\*<sub>GAS3</sub>-kanMX6* |  |
| RLY9878 | *trp1::P<sub>GPD</sub>-MTS-mCherry-GFP<sub>1-10</sub>-natMX6; LSG1-HA-GFP<sub>11</sub>-hisMX6; GAS1-GPI\*<sub>GAS5</sub>-kanMX6* |  |
| RLY9879 | *HSP104-GFP-his3MX6; trp1::P<sub>GPD</sub>-MTS-mCherry-natMX6; Δgas1-kanMX6-Leu2-P<sub>GAS1</sub>-gas1<sup>E161Q</sup>* |  |
| RLY9880 | *HO::P<sub>GAS1</sub>-gas1<sup>E161Q</sup>-GFP-hphMX6; trp1::P<sub>GPD</sub>-MTS-mCherry-natMX6* |  |
| RLY9881 | *trp1::P<sub>GPD</sub>-GRX5-HA-GFP<sub>1-10</sub>-natMX6; amp::GEM-P<sub>GAL1</sub>-FlucSM-HA-GFP<sub>11</sub>-kanMX6; TOM70-mCherry-Ura3MX6; Δgas1-his3MX6; kanMX6-Leu2-P<sub>GAS1</sub>-gas1<sup>E161Q</sup>* |  |
| RLY9882 | *trp1::P<sub>CUP1</sub>-Ub-R-EGFP-natMX6* |  |
| RLY9883 | *trp1::P<sub>CUP1</sub>-Ub-R-EGFP-natMX6; Δgas1-kanMX6* |  |
| RLY9884 | *Δbgl2-kanMX6* | 43 |
| RLY9885 | *Δgas5-kanMX6* | 43 |
| RLY9886 | *ura3Δ0::GEM-hphMX6; trp1::P<sub>GPD</sub>-GRX5-HA-GFP<sub>1-10</sub>-natMX6; HO::P<sub>GAL1</sub>-FlucSM-HA-GFP<sub>11</sub>-His3MX6; TOM70-mCherry-Ura3MX6* | 6 |
| RLY9887 | *ura3Δ0::GEM-hphMX6; trp1::P<sub>GPD</sub>-GRX5-HA-GFP<sub>1-10</sub>-natMX6; HO::P<sub>GAL1</sub>-FlucSM-HA-GFP<sub>11</sub>-His3MX6; TOM70-mCherry-Ura3MX6; Δbgl2-kanMX6* |  |
| RLY9888 | *ura3Δ0::GEM-hphMX6; trp1::P<sub>GPD</sub>-GRX5-HA-GFP<sub>1-10</sub>-natMX6; HO::P<sub>GAL1</sub>-FlucSM-HA-GFP<sub>11</sub>-His3MX6; TOM70-mCherry-Ura3MX6; Δgas5-kanMX6* |  |
| RLY9889 | *ura3Δ0::GEM-hphMX6; trp1::P<sub>GPD</sub>-GRX5-HA-GFP<sub>1-10</sub>-natMX6; HO::P<sub>GAL1</sub>-FlucSM-HA-GFP<sub>11</sub>-His3MX6; TOM70-mCherry-Ura3MX6; GAS1-GPI\*<sub>GAS3</sub>-kanMX6* |  |
| RLY9890 | *ura3Δ0::GEM-hphMX6; trp1::P<sub>GPD</sub>-GRX5-HA-GFP<sub>1-10</sub>-natMX6; HO::P<sub>GAL1</sub>-FlucSM-HA-GFP<sub>11</sub>-His3MX6; TOM70-mCherry-Ura3MX6; GAS1-GPI\*<sub>GAS5</sub>-kanMX6* |  |
| **Plasmid** | **Sequence** | **Reference** |
| RLB918 | *TRP1::P<sub>GPD</sub>-MTS-mCherry-natMX6* | 6 |
| RLB1051 | *GEM-P<sub>GAL1</sub>-FlucSM-HA-GFP<sub>11</sub>-KanMX6* | 7 |
| RLB1075 | *pRS316-HO(homology)-P<sub>GAS1</sub>-gas1<sup>N528K</sup>-GFP-hphMX6-HO(homology)* |  |
| RLB1076 | *pRS316-HO(homology)-P<sub>GAS1</sub>-mCherry-GAS1-hphMX6-HO(homology)* |  |
| RLB1077 | *pRS316-HO(homology)-P<sub>GAS1</sub>-GPI\*-GFP-hphMX6-HO(homology)* |  |
| RLB1078 | *pRS316-HO(homology)-P<sub>GAS1</sub>-SS-GPI\*-GFP-hphMX6-HO(homology)* |  |
| RLB1079 | *P<sub>GAS1</sub>-gas1<sup>E161Q</sup>-Leu2-KanMX6* |  |
| RLB1080 | *TRP1::P<sub>CUP1</sub>-Ub-R-EGFP-natMX6* |  |
| RLB1081 | *pRS316-HO(homology)-P<sub>GAS1</sub>-gas1<sup>E161Q</sup>-GFP-hphMX6-HO(homology)* |  |

To construct *gas1^N528K^* and *gas1^E161Q^*mutants with a GFP tag, Gas1-GFP was cloned into a plasmid, and underwent site-directed mutagenesis, before stably integrated into the *HO* locus. To construct *gas1^E161Q^* mutants for the spotting assay, split-GFP assay, and imaging of protein aggregates, a plasmid containing *gas1^E161Q^* was linearized and integrated into the *kanMX6* locus of the corresponding *Δgas1* strain. For the dually labeled mCherry-Gas1-GFP strain, mCherry-Gas1 and mCherry-Gas1-GFP was cloned and integrated into the *HO* locus of *Δgas1* strain. The last 31 residues of Gas1 after the cleavage site (GPI*) were fused with a GFP at the C terminus to construct the GPI*-GFP, and the ER-targeting signal peptide of Gas1 was added to the N terminus to construct the SS-GPI*-GFP. To replace the GPI* sequence of endogenous *GAS1*, the GPI anchor signal sequence of *GAS3* (last 26 residues) or *GAS5* (last 22 residues) together with a *kanMX6* selective cassette was amplified from corresponding plasmids in the Molecular Barcoded Yeast (MoBY) ORF Library (45) and inserted into the *GAS1* gene right after the sequence of N528 residue. Replacements were validated by colony PCR and sequencing. All variants of *GAS1* were expressed under *GAS1* promoter. Ub-R-EGFP under *CUP1* promoter was cloned from the plasmid pYES2-Ub-R-EGFP (Addgene #11953) (29) and stably integrated into the *TRP1* locus.

Standard YEP (yeast extract-peptone) supplemented with 2 % glucose (YPD) was used for transformations, biochemical analyses, and spotting assays. Synthetic complete (SC) supplemented with 2 % glucose was used for growing cells for confocal and super-resolution imaging. Yeast culture and plates were incubated at 30 °C except during heat shock at 42 °C. Optical density at 600 nm (OD_600_) was used to estimate the amount of yeast cells used in the various experiments.

### Drug treatments

β -estradiol (E2758, MilliporeSigma) was dissolved in ethanol and added to a final concentration of 1 μM for 90 min. Tunicamycin (T7765, MilliporeSigma) was dissolved in DMSO and used at a final concentration of 10 μg/ml for 2 h. DTT (R0861, Thermo Fisher) was dissolved in H_2_O and added into the yeast cell cultures at a final concentration of 10 mM for 2 h. Cycloheximide (239764, MilliporeSigma) was dissolved in DMSO and 100 μg/ml was used to treat cells for the indicated period. Calcofluor white (F3543, MilliporeSigma) was dissolved in DMSO to 10 mg/ml as stock and added in autoclaved YPD media plus 2 % agar at a final concentration of 10 μg/ml before agar solidification. CuSO_4_ (C1297, MilliporeSigma) was dissolved in H_2_O and 1 mM was used to treat cells for 30 min.

### Confocal microscopy and imaging conditions

Live cell images were acquired using a Yokogawa CSU-10 spinning disc on the side port of a Carl Zeiss 200 m inverted microscope. Laser 488 or 561 nm excitation was applied to excite GFP or mCherry, respectively, and the emission was collected through the appropriate filters onto a Hamamatsu C9100-13 EMCCD on the spinning disc confocal system. Regarding the multi-track acquisition, the configuration of alternating excitation was used to avoid the bleed-through of GFP. The spinning disc was equipped with a 100×1.45 NA Plan-Apochromat objective. Yeast culture condition for imaging: yeast cells were cultured in SC plus 2 % glucose overnight at 30 °C. The cells were then refreshed in the corresponding media for at least 3 hours at 30 °C until reaching an OD_600_ of about 0.3. For the estradiol-GEM inducible system (7,45), 1 μM of β−estradiol was added to the media for 90 min at 30 °C. All images in the same experiments were acquired with the same laser and exposure settings. For yeast 3D imaging, 0.5 μm step size for 6 μm in total in Z was applied. Image processing was performed using ImageJ software (NIH). For visualization purposes, images were scaled with bilinear interpolation and shown as the middle Z panel or maximum intensity projection (MIP) on Z for individual or merged fluorescent channels.

### Super resolution imaging

Structured illumination microscopy (SIM) images were acquired with a GE OMX-SR Super-Resolution Microscope 3D Structure Illumination (3D-SIM) equipped with high-sensitivity PCO sCMOS cameras. GFP and mCherry were excited with 488 nm and 568 nm lasers, respectively. The SIM images were reconstructed with the Softworx and aligned following the Applied Precision protocols. 3D rendering was performed with Imaris (Oxford Instruments Group).

### Split-GFP quantification

Split-GFP fluorescence from confocal images was quantified by using a custom Python code described before (6). In brief, mCherry and GFP intensities were summed along the Z axis, and then subjected to a random walk segmentation of the background and watershed segmentation of adjoining cells. For each cell, the mCherry channel was thresholded at 5% of maximal value to detect mitochondria, and median GFP intensity within mitochondria was calculated as the spGFP intensity per cell. Mean spGFP intensities of at least three biological repeats were used for the following analyses. Quantifications were shown either in absolute intensity values with an arbitrary unit or as normalized spGFP intensities to highlight the relative changes from the wild-type cells.

### Yeast whole cell lysis and immunoblots

Yeast cells in the indicated background were collected by centrifugation and snap frozen in liquid nitrogen for storage. Pellets were disrupted, boiled in 120 μl LDS sample buffer with 40 mM DTT (Thermo) for 10 min, and vortexed with an equal volume of 0.5 mm acid-washed glass beads to break cells at 4 °C for 2 min with 1 min interval. Cell lysates were re -boiled for 5 min, separated from glass beads by 15,000 *g* centrifugation at room temperature for 30 sec, and analyzed by SDS-PAGE.

Gel transfer was performed in iBlot2 (Thermo) and immunoblots were developed using Clarity Western ECL substrate (Bio-Rad) for HRP-linked secondary antibodies, or directly using fluorescent IRDye secondary antibodies (LI-COR). Data were acquired by using LI-COR imaging system and analyzed in Image Studio (LI-COR). HA-tag (C29F4) rabbit mAb #3724 from Cell Signaling Technology. PGK1 mouse mAb (22C5D8) from Invitrogen. GFP Living Colors A.v. mAb clone JL-8 (632381) from Takara Bio. Ubiquitin mouse mAb (P4D1) from Santa Cruz Biotechnology. Gas1 (N-terminal) antiserum was kindly provided by Dr. Hongyi Wu at the Mechanobiology Institute, NUS, Singapore.

### Mig1 nucleocytoplasmic translocation

Nucleocytoplasmic distribution of Mig1-GFP was quantified using a custom ImageJ macro and MATLAB script as published previously (46). Nuclear protein Pus1-RFP was used to create nucleoplasmic mask for the individual cell. Cytoplasm was defined by a dilated nuclear mask. The nuclear-cytoplasmic ratio of each cell was calculated by dividing the mean nuclear intensity by the mean cytoplasmic intensity. Populational means nuclear-cytoplasmic ratio of at least three biological replicates were used for statistical analyses.

### Yeast spotting assay

Single colonies of wild-type and mutant cells were inoculated in YPD media at 30 °C overnight. The cultures were diluted to the same OD_600_ of 1 and spotted at 10× serial dilutions from left to right on YPD plates containing 0.01% DMSO as control or 10 μg/ml CFW. Plates were then incubated at 30 °C for at least 2.5 days before scanning.

### Cycloheximide chase assay

Degradation of endogenous Lsg1 proteins after heat shock was evaluated as described previously (6). Briefly, a log-phase culture of yeast expressing Lsg1-HA was heated at 42 °C for 30 min. Recovery at 30 °C was performed in the presence of 100 μg/ml cycloheximide (CHX). At the indicated time points, the same volume of culture was collected, lysed by boiling and glass bead beating in LDS sample buffer, and subjected to immunoblotting analysis. To assess the UPS-mediated degradation, Ub-R-EGFP expression was induced in the presence of 1 mM CuSO_4_ for 30 min at 30 °C, followed by treatment with 100 μg/ml cycloheximide for 1 h. Single-cell GFP fluorescence was measured by using Attune NxT flow cytometer equipped with appropriate filter sets. Mean single-cell GFP intensities of three biological repeats were quantified and plotted.

### Hydropathy plots

Kyte-Doolittle hydropathy plots were generated using ProtScale (47,48) and the following parameters: window size, 5 amino acids; relative weight of the window edges compared to the window center, 100 %; linear weight variation model; no normalization. Transmembrane domain was predicted using TMHMM-2.0 (49).

### Statistical analysis

Descriptions of statistical tests and *P* values can be found in Figure Legends. Statistical analyses were performed with GraphPad Prism 6.0. No statistical methods were used to predetermine the sample size. The experiments were not randomized, and the investigators were not blinded to allocation during experiments and result assessment.

## Acknowledgments

We thank Dr. Hongyi Wu for providing Gas1 antiserum, and Drs. Susan Michaelis, Steven Claypool, Michael Matunis, and Jin Zhu for valuable discussions.

## Author Contributions

Conceptualization, investigation, visualization, and writing: YW, LR, RL Project supervision: RL

## Data and materials availability

All data needed to evaluate the conclusions are available in the manuscript. Additional data and materials related to this study can be requested from the authors. Requests for resources and reagents should be directed to and will be fulfilled by the corresponding author, RL.

---

**Supplementary Figure 1.**
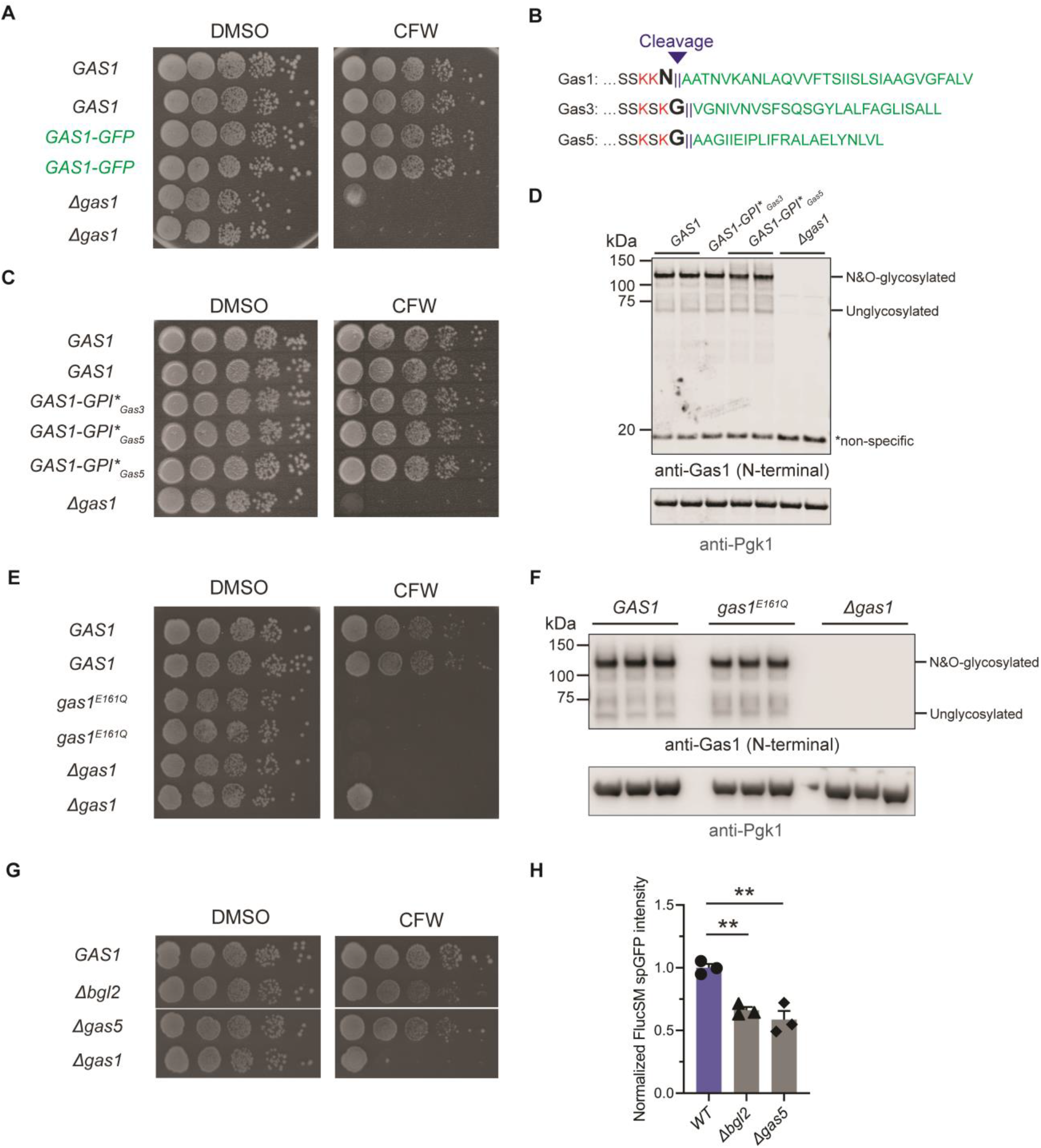
Characterization of Gas1-related mutants. **(A, C, E, G)** Yeast spotting assays on YPD plates containing DMSO or calcofluor white (CFW). Ten-fold serial dilutions from left to right for each plate. (**B**) C-terminal sequences of three Gas family proteins. Red, key basic residues; Bold, GPI-anchored attachment sites; Green, GPI anchor signals to be removed in matured proteins. (**D, F**) Immunoblots of whole cell lysates with wild-type, GPI* replaced, or catalytically inactive Gas1 using antiserum against the N-terminus of Gas1. (**H**) Quantification of mitochondrial FlucSM spGFP signal in wild-type cells and two cell wall mutants. Means ± SEM of normalized FlucSM spGFP intensities are shown. Unpaired two-tailed *t*-test. \*\**P* < 0.01.

**Supplementary Figure 2.**
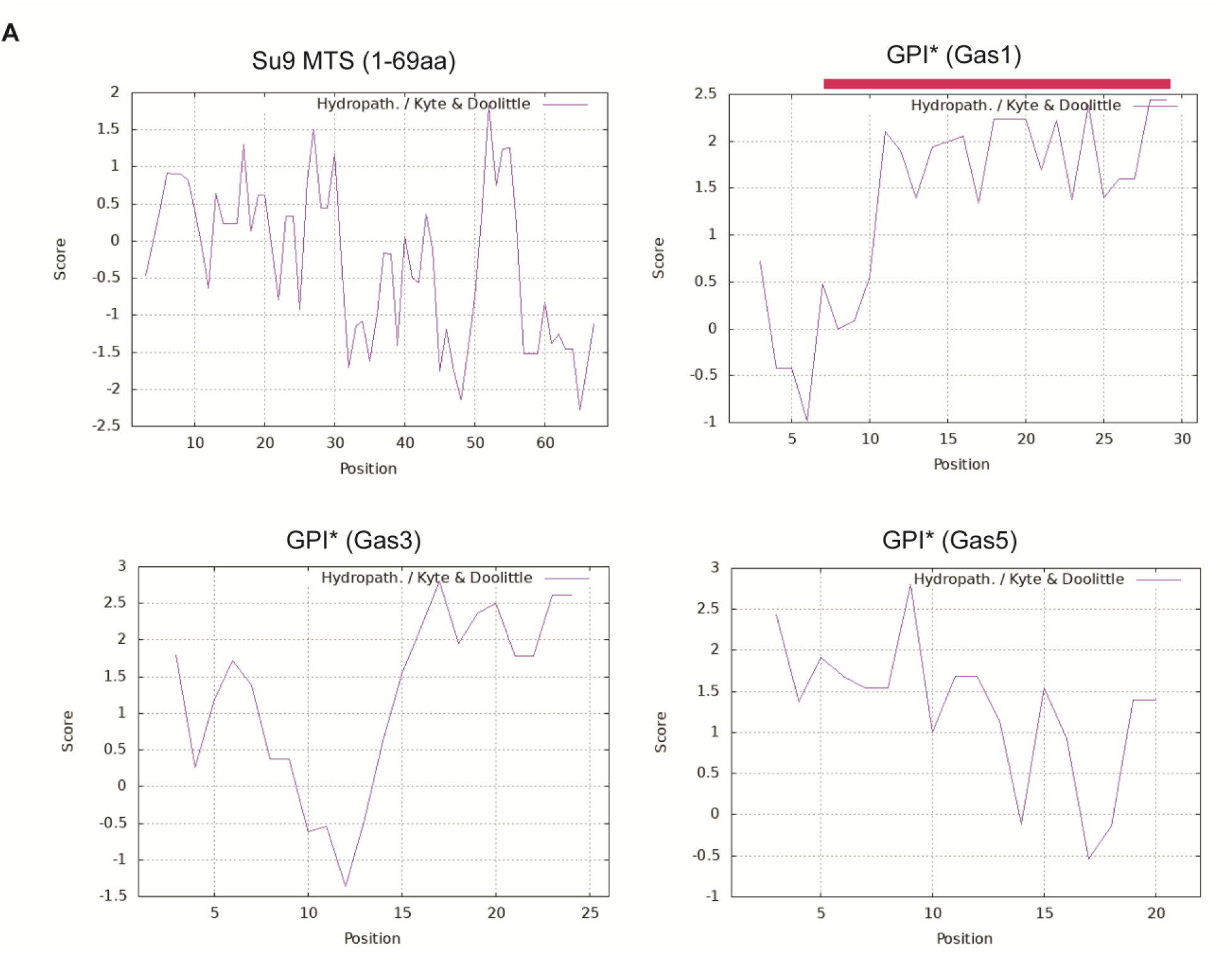
Sequence analyses of Su9 MTS and GPI anchor signals. (**A**) Kyte-Doolittle hydropathy plots generated by ProtScale. Su9 MTS is the first 69 residues of ATP synthase subunit 9 from *Neurospora crassa* (primary accession: P00842). GPI* of yeast Gas1, Gas3, and Gas5 represent the GPI anchor signal peptides. A transmembrane domain predicted by TMHMM-2.0 is highlighted as red bar.

